# Hearts may grow concentrically to balance ATP supply and demand and eccentrically to stabilize titin-based stress

**DOI:** 10.64898/2026.05.10.724147

**Authors:** John R. Kotter, Steve Leung, Thomas Kampourakis, Lik-Chuan Lee, Jonathan Wenk, Michael Moulton, Bertrand C. W. Tanner, Stuart G. Campbell, Christopher M. Yengo, Kerry S. McDonald, Julian E. Stelzer, Kenneth S. Campbell

**Author notes:** Corresponding author Kenneth S. Campbell, MS508 UKMC, 780 Rose Street, Lexington, KY 40536-0298.

## Abstract

Hearts change their wall thickness (concentric growth) and chamber size (eccentric growth) as they adapt to circulatory demands and the intrinsic function of their contractile cells. Factors associated with wall thickening include variants of sarcomeric proteins that enhance contractility, mitochondrial dysfunction, and hypertension. Chambers can dilate due to many factors including sarcomeric variants that depress contractility and aortic and / or mitral valve insufficiency. Despite intensive study, the mechanisms that regulate cardiac growth remain unclear. It is also uncertain whether inherited variants induce growth via the same mechanisms as more common clinical pathologies, such as hypertension. Here we show that computer simulations of a beating left ventricle reproduce both variant and non-variant-related growth patterns when myocytes grow concentrically to regulate intracellular ATP concentration and eccentrically to maintain titin-based intracellular stress. The simulations support the hypothesis that cardiac growth reflects homeostatic feedback through three interacting systems whereby myocytes add or remove mitochondria and sarcomeres (1) in parallel to match ATP generation to myocardial energy demand, and (2) in series to regulate passive tension, while (3) the autonomic nervous system regulates cardiac power, and thus myocardial ATPase, via baroreflex control. The new framework provides a mechanistic basis for the patterns of eccentric and concentric growth induced by a wide range of clinically-relevant conditions and could facilitate in silico testing of potential therapies for cardiac disease.

**Significance statement:** Hearts grow in response to both physiological and pathological stimuli. The patterns of concentric (wall thickening / thinning) and eccentric (chamber dilation / constriction) induced by different challenges are well recognized but the underlying mechanisms remain unclear. This work presents simulations of a beating left ventricle where (1) concentric growth is regulated by myocytes attempting to stabilize the intracellular ATP concentration and (2) eccentric growth is regulated by titin-mediated stress. The calculations reproduce the growth associated with inherited variants of sarcomeric proteins, mitochondrial dysfunction, hypertension, and both mitral and aortic valve insufficiency. The new ability to predict cardiac growth and its potential modification by treatments, including myotropes, brings the field closer to in silico optimization of therapy for cardiovascular disease.

## Introduction

One of the most important advances in molecular cardiology was the discovery that genetic variants in sarcomeric proteins can lead to familial cardiomyopathies (1, 2). The causal relationship was first established in the 1990s when variants of *MYH7*, the gene that encodes for cardiac β-myosin heavy chain, were linked to hypertrophic cardiomyopathy (3). Now, more than 20 sarcomeric genes have been linked to abnormal wall thickening (often called concentric growth) and associated familial hypertrophic disease (4). While the molecular mechanisms that lead to concentric remodeling remain unclear, most variants associated with hypertrophic disease produce a hypercontractile phenotype (5).

Other genetic variants lead to cardiac dilation and eccentric growth where the volume of the left ventricular chamber increases (6). Truncating variants in *TTN*, the gene that encodes for titin, are present in up to 60% of familial dilated cardiomyopathies and are the most frequent genetic cause of the condition (7). Variants in other sarcomeric proteins, including tropomyosin, troponin, and β-myosin heavy chain, can also induce dilation although this is less common (4).

In a pioneering study, Davis et al. combined experiments and computer modeling to investigate why variants induce distinct patterns of growth (8). Data from multiple mouse models of genetic cardiomyopathy showed that the area under an isometric twitch (the twitch-tension index) predicted whether hearts constricted and thickened or dilated and thinned. An important follow-up study built on this insight and showed that increasing the twitch-tension index by enhancing binding of Ca^2+^to the thin filament rescued the dilated phenotype of a mouse expressing a D230N variant of tropomyosin (9). Together, these papers reinforced the concept that variants that enhance contractility induce concentric growth while variants that reduce contractility produce wall thinning (10).

Excitingly, this science has had clinical impact. Mavacamten (11) and aficamten (12), drugs that suppress contractility by inhibiting myosin, are now approved for patients who have a specific type of hypertrophic cardiomyopathy where aortic outflow is partially blocked. Other sarcomere-acting drugs are under development and / or in trials (13). Patients who have inherited disease-inducing sarcomeric variants are likely to have new therapeutic options in coming years.

This progress should be celebrated but it is also important to remember that sarcomeric variants are not the most common cause of pathological growth in most clinical settings. At the University of Kentucky, a medium-sized academic medical center in the United States, 20,000 patients will undergo echocardiography each year. 2,500 of these individuals will show clinically significant wall thickening. Perhaps 100 will ultimately be diagnosed with a familial cardiomyopathy. 2,000 of the individuals with thick walls will have long-standing hypertension; 400 will have aortic stenosis. Sustained and substantial exercise will account for the wall thickening in a further handful of individuals. Similarly, most patients with a dilated left ventricle will not be diagnosed with genetic disease. At the University of Kentucky, eccentric growth due to sarcomeric variants is thought to occur ∼10-fold less frequently than dilation associated with mitral valve insufficiency following an infarction.

What controls cardiac growth when the twitch-tension index has not been perturbed by a sarcomeric variant? There is a spectrum of possibilities. At one extreme, variant and non-variant-induced growth reflect completely different mechanisms. At the other end of the spectrum, hearts grow concentrically and eccentrically according to a single set of pathways that can be perturbed by both variant and non-variant effects. The second hypothesis might provide a simpler way to think about long-term growth in patients and thus a potential opportunity to develop better therapies for diverse clinical conditions. This work used computer modeling to explore the possibility of a single unifying framework for cardiac growth.

The investigators started with three broad hypotheses. First, while experiments with mice had shown the power of Davis et al.’s twitch-tension index for predicting growth, stronger twitches involve more than just additional myosin cycling. It seemed possible that integrated force was a confounding variable that in variant-induced growth acted as a proxy for some related but currently unknown factor that also modulated growth in non-variant situations. Building on that idea, the authors noted that mitochondrial dysfunction can also induce concentric hypertrophy (14, 15). Moreover, mitochondrial protein content had been shown to increase when engineered heart tissues performed work by shortening against a load (16). Combining these concepts led to the hypothesis that the heart adapts its thickness to match ATP supply to energetic demand. Specifically, the authors envisaged a situation in which myocytes added or removed myofibrils and mitochondria in parallel until the rate of ATP generation matched the rate at which the ATP was consumed by the sarcomeres and ionic pumps. Precise identification of the underlying pathways was considered beyond scope for this work but Davis et al. suggested that ERK1/2 could be involved (8, 17).

Second, the authors recognized that human ventricular walls do not always thin as a patient’s chamber dilates. This contrasts with the behavior observed in mice with sarcomeric variants and led to the hypothesis that eccentric growth, where sarcomeres and mitochondria are added to myocytes in series, is regulated by a different mechanism than concentric growth. In skeletal muscle, eccentric growth was recently shown to be sensitive to titin (18). It thus seemed appropriate to test a similar mechanism in hearts whereby myocytes add or remove sarcomeres and mitochondria in series to provide negative feedback control of titin-based stress.

Third, the amount of ATP used by the heart depends on the rate at which blood is pumped. Since ATP consumption was hypothesized to influence concentric growth, cardiac power would need to be regulated as occurs in vivo through the action of the autonomic nervous system. Hearts that were generating more power, as occurs, for example, during hypertension or valvular disease, would need to be bigger.

This manuscript describes simulations that integrated these three concepts. The calculations were performed using FiberVent, which extended the previously published MyoVent (19, 20) by replacing the cross-bridge distribution based calculation of sarcomeric force (21) with the spatially-explicit FiberSim framework (22). This allowed the new calculations to test how different hypothesized actions of myosin binding protein-C would influence cardiac growth. Arterial pressure was regulated by an in silico baroreflex that continually adjusted heartrate, excitation-contraction coupling, thick and thin filament regulation, and vascular tone to stabilize arterial pressure (19).

FiberVent is open-source (23). The code repository includes instructions that reproduce all results and figures presented in this manuscript.

## Results

### Simulated ventricles grew to a steady-state

Simulations were performed using the FiberVent framework that is described in the Methods section. Figure 1 shows a subset of the signals simulated by the model for four cardiac cycles. The baroreflex had stabilized mean arterial pressure at a target setpoint of 90 mmHg but the growth algorithms had not been activated. The wall was thicker in this starting configuration than required to maintain cardiac output so the intracellular ATP concentration trended upwards as the mitochondria produced more ATP than the myocytes used to maintain arterial pressure.

**Figure 1.**
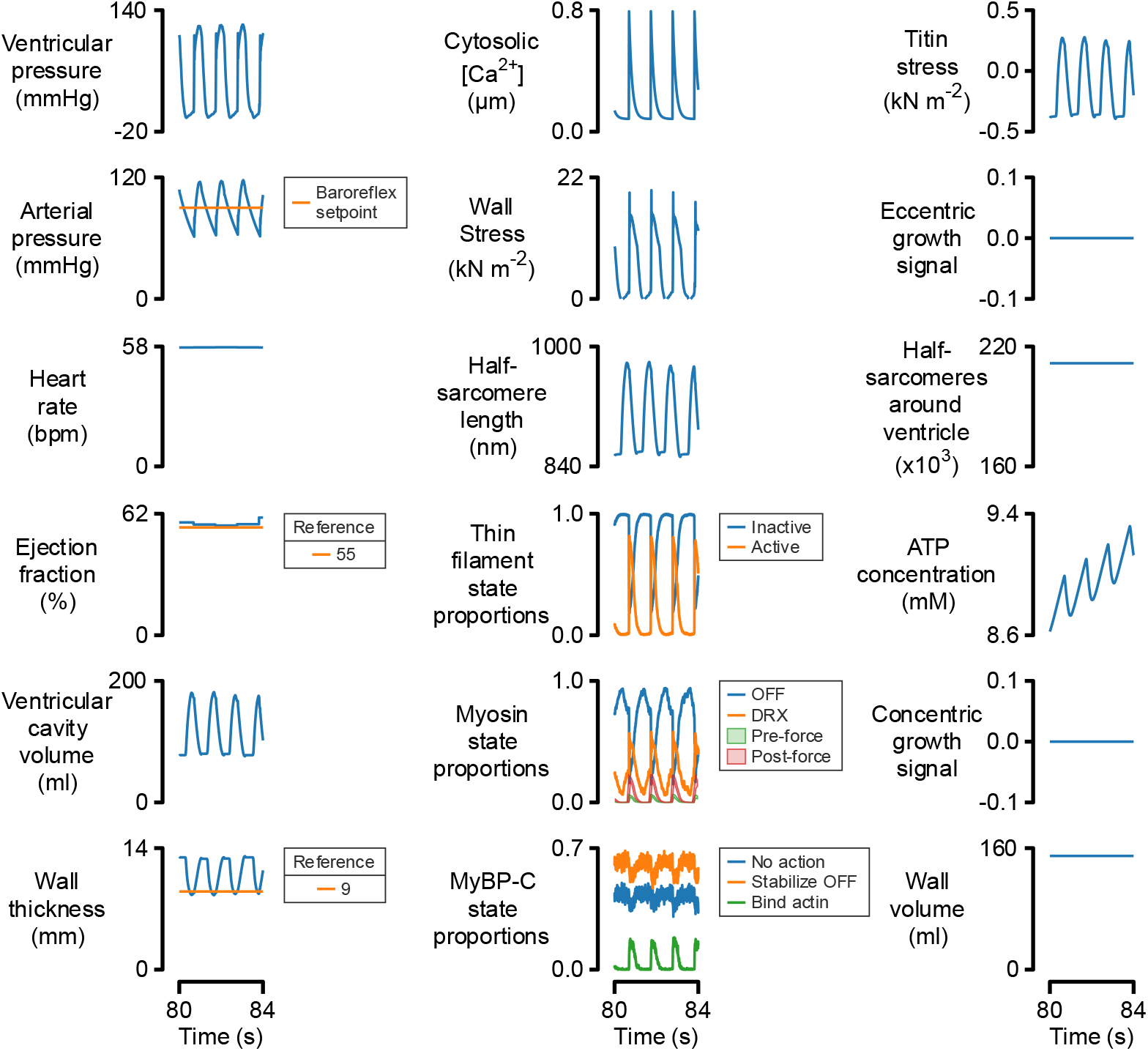
Cardiovascular function for four beats without growth. Traces were simulated using a version of the default model where the wall thickness and the number of circumferential half-sarcomeres had been perturbed from their values after growth. The baroreflex had stabilized arterial pressure at the target of 90 mmHg but the growth algorithms had not been activated. Note that the intra-myocyte ATP concentration (middle of right-hand column) cycles with each contraction but trends upwards with time indicating that the mitochondria are generating ATP more quickly than the myocytes are consuming it.

Figure 2 shows how the system responded when the growth algorithms were activated. The transients represent interactions between the feedback systems. Note that before growth, the intra-myocyte ATP concentration was rising over time (fourth row, right-hand column) while titin-based stress was below its setpoint value (top row, right-hand column). This implies that the ventricle’s myocytes consumed less ATP than their mitochondria were generating and that the myocytes were longer than required to create the homeostatic value of titin-based stress.

**Figure 2.**
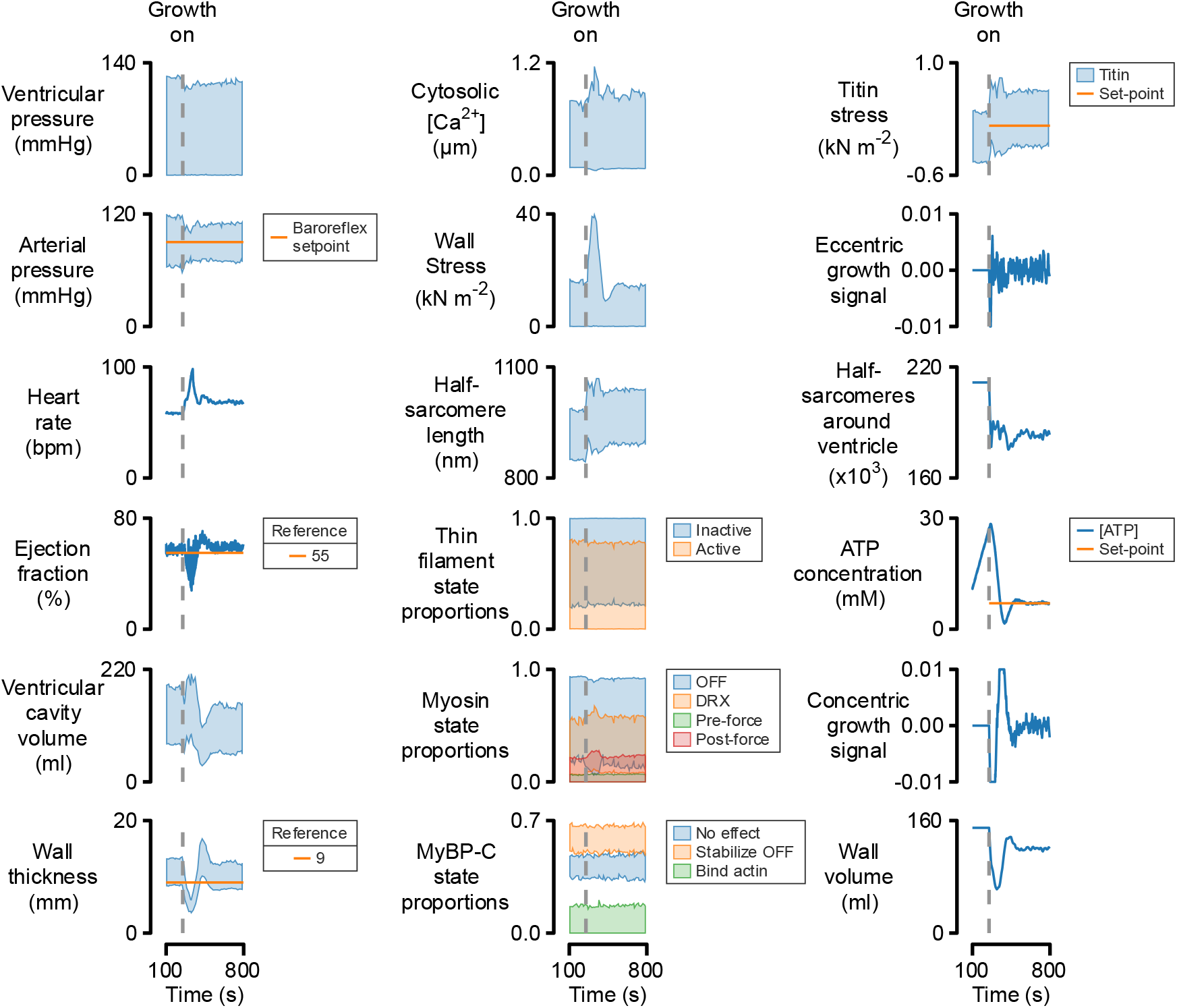
Cardiovascular function as the heart grew to balance energy supply and demand and stabilize titin-based stress. Simulations as described for Figure 1. The growth algorithms were activated at 250 s after which the ventricle grew concentrically and eccentrically until the intracellular ATP concentration, titin-related forces, and arterial pressure were maintained at their respective setpoints by the interacting feedback systems. The system was stable in this configuration and would remain there indefinitely unless it was perturbed. Signals that varied markedly during each cardiac cycle (for example, ventricular pressure) are shown as envelopes that indicate the extreme values. Overlap of the transparent envelopes produces five color shades for the myosin state populations.

When the ventricle grew, the heart removed sarcomeres and mitochondria in series (third row, right-hand column) to drive titin to its set-point and adjusted wall thickness (bottom row, left-hand column) to better match ATP supply to demand. Over time, the ventricle evolved to a new configuration where ATP supply was balanced with energetic demand, the time-average of titin-based stress was controlled at its setpoint, and arterial pressure was at target. The ventricle had grown slightly thinner and had a lower end-diastolic volume (fifth row, left-hand column). Since the system had reached steady-state, it would remain stable in this configuration until perturbed.

### Increased contractility induces wall thickening and chamber constriction

One of the advantages of computer modeling is that parameters can be adjusted to test hypotheses. Figure S1 in Supplementary Material shows how the system evolved when myosin-binding protein-C molecules become less effective at stabilizing myosin molecules in their OFF / super-relaxed / interacting heads motif configuration (24). The perturbation allowed more myosin heads to cycle during systole so the ventricle used ATP more quickly. This caused the myocytes to add myofibrils and mitochondria in parallel to distribute the increased energetic demand over a greater number of mitochondria. The thickening associated with this concentric growth restricted diastolic filling so titin-based forces dropped. The myocytes reacted by reducing the number of sarcomeres in series to return titin-based stress to its target level. The simulated ventricle grew to become thicker with a smaller chamber volume, consistent with findings in some patients who inherit variants of myosin-binding protein-C.

Figure 3 summarizes the growth patterns predicted by multiple simulations testing different interventions. The filled circles in Figure 3A show how the ventricle grew when model parameters associated with myofilament function or intracellular calcium handling were adjusted. The color-coding shows the rate of ventricular pressure development during isovolumic contraction. Perturbations with higher rates of pressure development (and thus increased contractility) produced smaller, thicker ventricles. These model-based predictions reproduce Davis et al.’s experimental results from genetically-manipulated mice with sarcomeric variants (8).

**Figure 3.**
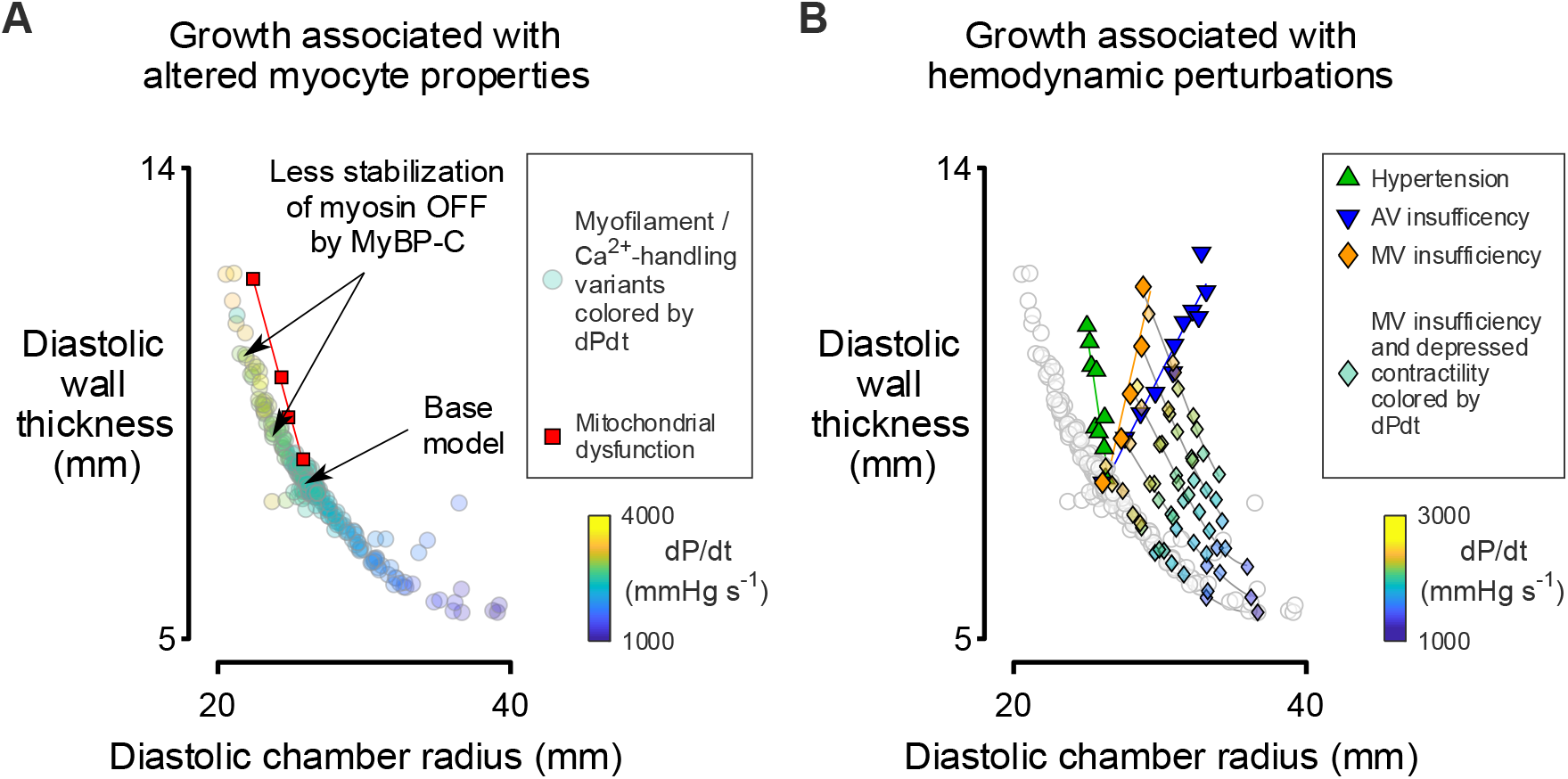
Predictions for growth induced by altered myocyte properties and perturbations to system-level hemodynamics. (A) Circles summarize ventricles where myofilament or calcium-handling related properties were adjusted to change contractility. Symbols are color-coded by the rate of pressure development during isovolumic contraction. Squares show growth for different levels of mitochondrial dysfunction (5 to 20% reduction in ATP production). (B) Upwards-pointing triangles show simulations where the setpoint of the baroreflex was increased to mimic hypertension (maximum mean arterial pressure of 120 mmHg). Downwards-pointing triangles show the response to aortic valve insufficiency. Diamonds show growth induced by increasing mitral valve insufficiency with lines sloping down to the right indicating further dilation when contractility was reduced via perturbed myofilament contractility.

### Mitochondrial dysfunction induces concentric growth

Mitochondrial dysfunction can also induce concentric hypertrophy (14, 15). This effect was mimicked by the squares in Figure 3A which show the effect of successively reducing the rate at which the mitochondria generated ATP. The walls thickened in the simulations as the myocytes added mitochondria to sustain the energetic demand of maintaining arterial pressure. The thicker walls restricted filling so sarcomeres were removed in series via eccentric growth pathways to restore titin-based forces. This led to chamber constriction. Since the perturbations to mitochondrial function did not alter the intrinsic contractility of the myocytes, ventricles grew thicker for a given chamber diameter (squares in Figure 3A) than predicted for tests modulating sarcomere or excitation-contraction related function (circles).

### Increased energetic demand induces concentric growth

Figure 3B shows the effect of changing energetic demand. The upwards-pointing triangles demonstrate how the ventricles grew as the set-point for the baroreflex was progressively raised to mimic systemic hypertension. Maintaining the increased pressure required greater cardiac power so the myocytes grew thicker by adding myofibrils and mitochondria in parallel to distribute the workload across more tissue. As described above, the increased thickness induced chamber constriction, again resulting in a thicker but slightly smaller ventricle.

Similar effects were observed when the arteriolar resistance was reduced to mimic one of the physiological responses to aerobic exercise. As shown in Figure S3 and S4, the baroreflex increased heart rate, contractility, and venous tone to maintain arterial pressure and the ventricle thickened to compensate. Although preliminary, these observations suggest that the modeling framework can predict some of the structural changes associated with sustained exercise (25).

### Valvular insufficiency induces chamber dilation and wall thickening

As shown in Figure S5, aortic insufficiency is a clinical condition where blood leaks backwards into the ventricle during diastole. The downwards-pointing triangles in Figure 3B show that the simulations reproduced the well-known clinical phenotype associated with this condition where the ventricle thickens as it dilates. In the simulations, the growth response reflected the increased diastolic stretch induced by the regurgitant flow. Since the aortic pressure exceeded the typical venous filling pressure, titin molecules were stretched further than normal, thereby activating eccentric growth and serial addition of sarcomeres and mitochondria. Further, since blood was leaking backwards from the aorta, arterial pressure dropped below at its target setpoint. This temporary reduction was corrected by the baroreflex which adjusted contractility, heart rate, and vascular tone to increase forward flow and return arterial pressure to its setpoint. The compensation required additional power so the ventricular wall thickened to match mitochondrial ATP production to the increased demand (Figure S6).

Mitral valve insufficiency produced a similar cascade of feedback-linked responses but the ventricles dilated less for a given increase in wall thickness (orange diamonds in Figure 3B) than during aortic insufficiency. As explained above, the wall thickening reflected the increased myocardial power needed to sustain arterial pressure in the presence of regurgitation. However, at least in the current simulations, the pressures filling the ventricle were lower in the presence of mitral regurgitation than they were during aortic insufficiency. This reduced the excess strain of titin molecules and diminished the eccentric growth response.

### Reduced contractility exacerbates chamber dilation in the presence of valvular insufficiency

Patients with severe cardiac disease rarely have just one problem. For example, many individuals develop mitral valve insufficiency following an infarction that reduces myocardial contractility. When mitral valve insufficiency and reduced contractility were tested simultaneously (color-coded diamonds in Figure 3B), FiberVent predicted that they produced additive effects. Note in Figure 3B how the reduced contractility induced progressive dilation and wall thinning, similar to the effects predicted in the absence of valvular disease (circles, Figure 3A).

## Discussion

### A single mechanistic framework can predict realistic growth responses to different stimuli

These simulations demonstrate that a single mechanistic framework can reproduce the different patterns of concentric and eccentric growth induced by perturbations to the function of sarcomeres, mitochondria, and / or calcium-handling systems and by modifications to system-level hemodynamics. The calculations do not prove that hearts grow to stabilize the intracellular ATP concentration and titin-based stress but they do show that growth patterns seen in genetically-modified mice and in patients with different forms of cardiac disease are consistent with feedback based on these signals. Moreover, the simulations align with experiments which show that mitochondrial protein levels increase with contractile work when engineered heart tissues shorten against different loads (16). Future experiments, perhaps involving engineered heart tissues stimulated at different rates and shortening from different lengths, could be used to test the growth laws experimentally.

The new ability to predict how ventricles adapt their diameter and thickness as they remodel provides an opportunity to further integrate computer models into the development of better therapies for cardiac disease (26). This could be particularly important for the further development of myotropes (13) because newly enhanced computer modeling could reduce the need for animal-based testing by predicting how a patient’s heart will adapt over weeks and months of treatment.

### Growth associated with protein variants reflects dynamic interactions between feedback loops

As described in Methods and Materials, the FiberVent framework integrates a recognized model of myofilament biophysics into a system-level model of the circulation. Arterial pressure is regulated by a feedback loop that mimics some of the physiological actions of the baroreflex system. Further feedback loops modulate parallel and series addition of sarcomeres and mitochondria in response to changes in the intracellular concentration of ATP and titin-based stress respectively. Concentric and eccentric growth reflect dynamic interactions between these systems.

The growth patterns predicted by hypothetical variants of myosin binding protein-C provide a practical example. Variants that are less effective at stabilizing myosin heads in their OFF configuration augment contractility, and thus ejection fraction, by allowing more heads to interact with actin during systole. This allows the heart to maintain cardiac output, and thus the arterial pressure (which is being maintained via autonomic control), with a smaller chamber. Since more cross-bridges are cycling, the sarcomeres are also consuming ATP at a faster rate. The myocytes thicken by adding mitochondria and myofibrils in parallel so that the contractile work required to maintain arterial pressure is shared over a greater volume of myocardium. In essence, the heart is growing concentrically to preserve the number of cycling heads supplied by a given volume of mitochondria. The result is a ventricle that is thicker and with a smaller chamber diameter. Two examples are indicated in the upper left quadrant of Figure 3A.

Variants of MyBP-C that suppress contractility induce the opposite effects. Hearts expressing these variants have a lower ejection fraction and need a larger end-diastolic diameter to maintain cardiac output. Power consumption at the sarcomere level is also reduced so the heart thins as it dilates. Examples of these situations are found at the lower right of Figure 3A but were not specifically labeled as the figure is already complex.

These general mechanisms are not specific to myosin binding protein-C. Any modification that perturbs myofilament contraction or calcium-handling will induce compensatory growth. Variants that augment contractility can maintain arterial blood pressure with smaller ventricles while loss of function implies a lower ejection fraction and thus a larger chamber. The circles in Figure 3A demonstrate the relationship with the color-coding showing higher contractility (quantified as the rate of isovolumic pressure development) inducing thicker ventricles with smaller chambers.

Since the ventricle changes thickness until its energy demand is matched by its energy supply, changes to mitochondrial function also induce growth. The squares in Figure 3A show the ventricular walls thicken as the rate at which mitochondria generate ATP falls. This result mimics the concentric hypertrophy observed in animals and patients with mitochondrial dysfunction (14, 15).

### Growth associated with perturbed hemodynamics reflects altered cardiac power

Hypertension and valve disease are the most common causes of pathological cardiac growth in most clinical settings. The current framework attributes growth in these situations to the ventricle’s increased use of energy.

The upwards-pointing triangles in Figure 3B show how the ventricle thickens as the set-point for arterial pressure in the baroreflex control system is progressively increased. Since the myocytes require more power to sustain the increased arterial pressure, more mitochondria are required to maintain the intra-myocyte ATP concentration at its target level. This induces the wall thickening and mild constriction that characterize long-term hypertension in patients.

Aortic insufficiency (summarized by the downwards-pointing triangles in Figure 3B) results in a different pattern of growth where the ventricle dilates as it thickens. As shown in Figure S5, aortic insufficiency is a condition where blood leaks backwards from the aorta into the ventricle during diastole. To maintain arterial pressure, cardiac output and thus power must increase to regain appropriate forward flow. As described for hypertension above, the increased demand for energy drives wall thickening. The additional challenge of aortic insufficiency is that the ventricle, and thus titin molecules within its constituent myocytes, experience an additional stretch from the regurgitant flow. Since the pressure gradient across the leaking valve is so high, the increased passive force is sufficient to induce marked eccentric growth.

### Comparison with the twitch-tension index explanation for growth

Much of this work was inspired by Davis et al.’s demonstration that the twitch-tension index (the area under an isometric twitch) predicts whether ventricles grow concentrically or eccentrically in mice when intrinsic contractility has been manipulated via genetic techniques (8). Since Davis et al.’s discoveries were a breakthrough for the field, it is important to clarify how the current work provides additional advances.

First, this work proposes that wall thickening and chamber dilation are regulated by different systems. In Davis et al.’s framework, ventricular wall thickening was always accompanied by chamber constriction implying a fixed coupling between concentric and eccentric growth. This is entirely appropriate when intrinsic contractility is the prime experimental variable – indeed, the circles in Figure 3A show the same relationship. However, fixed coupling limits application to clinical conditions, such as hypertension and valvular disease, where the heart faces additional hemodynamic challenges.

Second, contractility was not fixed in this work but was dynamically regulated through baroreflex control of arterial blood pressure. This is critical for accurate predictions. As an example, if the baroreflex is not activated, the simulations predict that aortic insufficiency induces chamber dilation with wall thinning rather than the wall thickening that occurs in real patients. This is because the regurgitant flow still stretches the ventricle but, without the baroreflex, there is no compensatory increase in cardiac power.

Third, this work provides a novel explanation for hypertrophy associated with variants of myosin binding protein-C. Moreover, the simulations provide opportunities to probe the underlying mechanisms. The basic premise of the growth shown in Figure S1 is that myocyte efficiency is reduced (that is, the cells consume more ATP for a given amount of cardiac work) when myosin binding protein-C is less able to stabilize myosin in a suppressed state. Still not clear though, is where this energy is lost. Potential energy sinks include the additional cycling heads, cooperative activation induced by myosin binding protein-C molecules attaching to the thin filament, and / or the need to overcome mechanical drag induced by these bound molecules. Each of these mechanisms can be analyzed in more detailed simulations to identify high value therapeutic targets.

In summary, the current simulations advance the field by linking a detailed biophysical model of myofilament function with growth algorithms and autonomic cardiovascular control. The cross-system integration may help to accelerate future applications to clinical settings.

### Predictions that could be tested in future experiments

Models can be used to predict results that can be tested in future experiments. Two of many potential examples are discussed below.

#### Prediction 1

Cardiac dilation in advanced heart failure is not induced by fibrosis.

The hearts of most patients who receive a cardiac transplant have dilated chambers, reduced contractility, and exhibit increased fibrosis (27, 28). In principle, the enhanced fibrosis could reduce wall stress (and thus contractility) by occupying space that would otherwise be filled by myofilaments. It could also add passive stiffness and induce dilation by diminishing the passive force born by titin molecules during diastolic filling. Both effects will induce dilation in the current FiberVent framework.

In practice, when the proportion of the ventricular wall occupied by fibrosis was increased in test simulations, the ventricle thickened and constricted (Figure S7). This potentially unexpected result can be explained by two factors. First, the total cardiac power required to sustain arterial pressure had not changed so the myocytes that were left in the myocardial wall were required to do more work. This induced concentric growth and wall thickening. Second, at least in the base-line model used throughout this work, the collagen matrix was relatively compliant over the normal operating range. (Note that this assumption is consistent with the observation that titin molecules bear most of the passive tension at physiological sarcomere lengths (29).) Consequently, titin-loading was only minimally affected by the increased volume of extracellular matrix and the effect on eccentric growth was outweighed by the increase in wall thickness required to maintain cardiac power.

#### Prediction 2

Increasing sarcomeric efficiency will reduce wall thickness.

Ventricular walls thicken or thin in the current simulations until the myocytes’ demand for ATP is balanced by the ability of their mitochondria to supply it. The heart’s power consumption is determined by the arterial pressure and the properties of the valves and blood vessels and is regulated through autonomic control.

One way to thin the wall is to help the mitrochondria generate ATP at a faster rate, as was done in a phase 2 trial testing ninerafaxstat (30). This novel mitotrope shifts mitochdondrial function towards glucose oxidation (31). Another strategy is to reduce the sarcomeres’ need for ATP. Since the cells will still be required to generate the same power, this implies an increase in efficiency. This aspect of myocardial function has likely been targeted by evolution but it might be possible to modulate non-optimal situations associated with disease.

As an example, note that initiating contraction with calcium is both quick and effective but comes at the cost of an indirect relationship between activation and demand. Thick filament mechano-sensing, where myosin heads are recruited directly by load (24), may be harder to initiate but ties energy usage more closely to need. In situations like heart failure, where energy supply is particularly challenging, judicious dosing with a calcium desensitizer might bias the system towards cooperative myofilament mechanisms that reduce unnecessary activity.

### Limitations

The authors acknowledge many limitations of this work but, for conciseness, will discuss just three.

Mitochondrial function is radically simplified in the current framework. The organelles are merely assumed to generate a defined number of ATP molecules per unit volume per second. There is a clear opportunity to enhance the simulations with a more sophisticated framework for mitochondrial function in future work (32).

FiberVent uses a ‘one-fiber’ approach to simulating ventricular function (33). This simplifies the calculations, but it eliminates the possibility of investigating regional growth effects including those associated with apical or septal-dominant hypertrophy. It would be exciting to implement region-specific growth in finite element calculations and test whether localized strain patterns impose distinct energetic demands and thus variation in growth (34). Although some way in the future, it might even be possible to work backwards from imaging data describing an abnormal cardiac structure to a molecular cause. Whether this biophysics-forward approach would be better than AI-based analyses of large clinical datasets remains open to question.

Even the corresponding author (trained as a biophysicist) recognized that FiberVent underestimates the number of chambers in a human heart. Implementing growth in a 4-chamber system that is appropriately coupled to pulmonary and systemic circulations is an obvious next step. This could be achieved via a simplified framework like CircAdapt (35), with a low-order model (36), or using full finite element approaches (37).

One problem in the current work is that during simulations of mitral valve insufficiency, blood was pumped backwards into the venous system rather than into the left atrium. Since the venous system had high compliance, filling pressure in the simulations did not rise to the levels measured in patients with severe disease. This explains why the ventricles exhibiting mitral valve insufficiency did not dilate as far in the simulations as those with aortic insufficiency (Figure 3B). While similar trends are seen in patients (38), the current simulations probably overestimated the real effect.

## Conclusions

Figure 4 summarizes the hypotheses tested in this work. Concentric growth reflects myocytes adding myofibrils and mitochondria in parallel to match ATP supply to energetic demand. Myocytes add sarcomeres and mitochondria in series to provide negative feedback control of titin-mediated forces. Cardiac output, and thus the demand for ATP, is regulated by the autonomic nervous system through baroreflex control.

**Figure 4.**
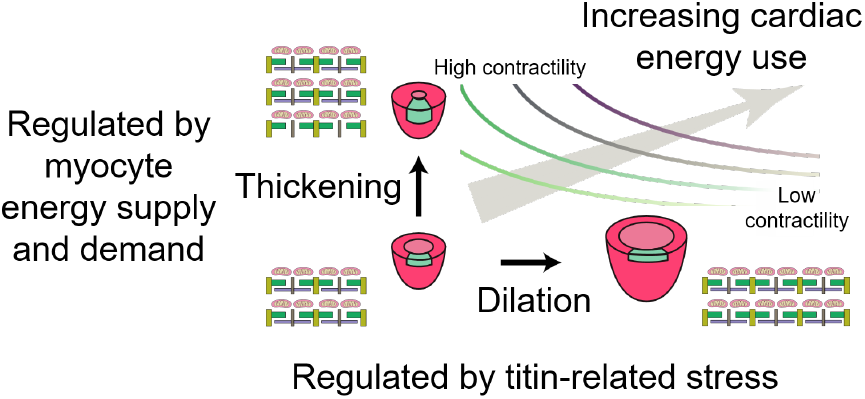
Framework for cardiac growth. This work tested the hypotheses that cardiac growth reflects homeostatic feedback through three interacting systems. Myocytes added or removed mitochondria and myofibrils in parallel to match ATP generation to energy demand, and in series to regulate passive tension. These processes were influenced by the autonomic nervous system which modulated contractility, and thus myocardial ATPase, to adjust cardiac output and maintain arterial pressure. The curved lines are contours showing ventricles of different sizes that consume ATP at the same rate.

Perturbations to cellular-level contractility, including those associated with inherited variants, induce growth along contours associated with equal cardiac energy use. Ventricles with high ejection fractions can maintain arterial pressure with a small chamber volume and traverse along the contour in the direction of wall thickening. Variants with lower contractility require a larger chamber to maintain arterial pressure and dilate along the same contour. Changes to the vasculature, arterial pressure, or valvular function change the ventricle’s demand for energy and induce growth to a different contour.

Working in consort, these interacting feedback systems predict realistic growth responses to a wide range of physiological and pathological stimuli. The current work enhances mechanistic understanding of cardiac growth and may help in the development of computer-assisted optimization of therapy for patients with cardiac disease (26).

## Methods

### Overview

The FiberVent model used for this work extended the published PyMyoVent framework (19) with an enhanced computational model of myofilament function, calculations pertaining to the intra-myocyte ATP concentration, and with growth algorithms. All are described below. The baroreflex controlling arterial pressure has been described in detail (19). The code is open-source and available via GitHub (23). The repository includes instructions to duplicate all the results included in this manuscript.

### Myofilament function

PyMyoVent simulated sarcomere-level function using cross-bridge distribution techniques (21). In FiberVent, the contractile module was replaced with the spatially-explicit FiberSim model of myofilament function (22). This provided the ability to predict how different modes of myosin binding protein-C function modulated growth.

As previously described (22), myosin heads were arranged in dimers and transitioned between (a) an inhibited OFF state that could not interact with actin, (b) an ON state (that could potentially attach to available binding sites on actin), (c) a pre-force-step actin-bound state, and (d) a post-force-step actin-bound state. Myosin binding protein-C molecules were positioned in the C-zone of each thick filament with triplets repeating every third myosin crown for a total of 9 stripes. Each myosin binding protein-C molecule cycled independently between a null state (where it had no physical function), an actin-bound state (where it contributed to cooperative activation of the thin filament and imposed drag), and a myosin stabilizing state (where it reduced the probability of neighboring myosin heads exiting the OFF configuration).

### Intra-myocyte ATP concentration

Calculations of the intra-myocyte ATP concentration assumed that the myocardial wall was formed from an extracellular matrix component (which occupied space and contributed to passive stiffness but was otherwise inert) and myocytes which, in turn, contained myofibrils and mitochondria. The myofibrils and ionic pumps consumed ATP while the mitochondria generated it.

If more ATP was consumed than generated, the ATP concentration rose over time. Conversely, if the myocytes used less ATP than their mitochondria generated, the intracellular ATP concentration declined.

These concepts were formalized as equation 1 which defined the rate of change of the intracellular ATP concentration as:

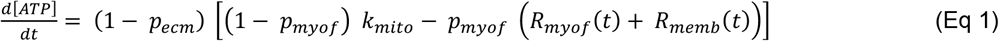

where *p*_*ecm*_ was the proportion of the wall occupied by extracellular matrix, *p*_*myof*_ was the proportion of myocytes occupied by myofibrils, *k*_*mito*_ was the rate at which a given volume of mitochondria generated ATP, and *R*_*myof*_*(t)* and *R*_*memb*_*(t)* were the rates at which the myofibrils and membranes consumed ATP respectively. The myofibrillar ATPase was determined from the rate at which myosin heads detached from the post-force-step actin bound state. Since the sarcoplasmic endoplasmic reticular calcium-ATPase (SERCA) uses most of the energy associated with maintenance of ionic concentrations in the heart (39), membrane-associated ATPase was approximated as:

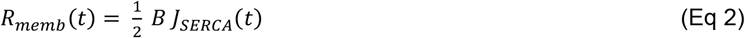

where *J*_*SERCA*_*(t)* represented Ca^2+^ions pumped by SERCA in the model, B was a scaling factor that accounted for the myocytes’ buffering capacity and the additional Ca^2+^ions that would need to be pumped in vivo, and the ½ term reflected the 2 Ca^2+^ions pumped by SERCA for each ATP molecule.

### Growth algorithms

FiberVent simulated growth by allowing the ventricle to change wall thickness and chamber diameter.

Concentric growth (changes in wall thickness) depended on the intracellular ATP concentration described in the preceding section. Eccentric growth (changes in chamber diameter) was controlled by the stress in titin molecules. The speed of growth was set by rate constants defined for each simulation. Further details, including the relevant equations, are described below.

Changes in wall thickness (*T*) mimic myocytes adding / removing myofibrils and mitochondria in parallel and were controlled by the intracellular ATP concentration via:

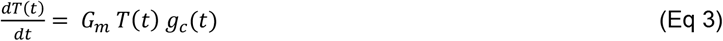

where G_*m*_ was the master growth rate and *g*_*c*_*(t)* was the concentric growth signal. Its rate of change was defined as:

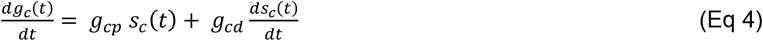

where:

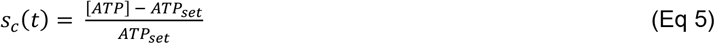

and *g*_*cp*_ and *g*_*cd*_ were scaling factors for proportional and derivative feedback and *ATP*_*set*_ was the homeostatic setpoint for the ATP concentration. Both scaling factors were held negative so that the wall thinned if the intracellular ATP concentration was above its setpoint. The derivative in equation 4 improved homeostatic control because altering wall thickness changed the rate at which the myocytes used ATP rather than the ATP concentration itself. Derivative control is common in human-created feedback systems and can be generated via molecular pathways (40).

The algorithms updated wall thickness without altering chamber diameter. This mimics symmetric transmural growth as opposed to myofibrils and mitochondria being added only to the endocardial or epicardial edges of myocytes.

Eccentric growth was simulated by varying the number of half-sarcomeres in series around the ventricular circumference (*n*) based on the stress in titin molecules (*F*_*titin*_). This mimics myocytes adding / removing sarcomeres and mitochondria in series. The relevant equations were:

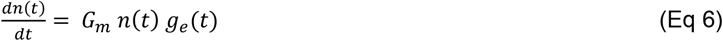

where:

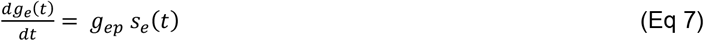

and:

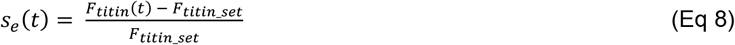

Derivative control was not required for eccentric growth because titin-related forces were inversely proportional to the number of half-sarcomeres around the heart’s circumference.

Growth in real hearts takes weeks to months which would be challenging to simulate. Fortunately, growth could be accelerated in the simulations without affecting the ventricle’s final dimensions. Figure S2 in Supplementary Material shows that as long as growth occurred slower than the baroreflex attempted to change cardiac function, the diameter and thickness of the heart at steady-state was not influenced by the master growth rate *G*_*m*_.

### Implementation and computer code

Based on experience with FiberSim, FiberVent was created as two components. The first, FiberVentCpp, was the core simulation code. It was designed to run as quickly as quickly as possible and was thus written in C++ with non-trivial calculations performed using the GSL scientific computing library (41). The C++ code has ∼14,000 lines and was compiled for standard Windows computers. The second component, FiberVentPy, consists of ∼6,000 lines of Python and provided an interface that simplified testing different conditions and analyzing simulation outputs.

Since the FiberSim contractile model used stochastic techniques, results needed to be averaged over multiple filaments to predict ventricular wall stresses. This took time. As an example, Figure 2 shows records calculated using a FiberSim model with 36 thick filaments. The complete simulation required ∼5 hours on a single PC thread. The ∼500 simulations summarized in Figure 3 were completed in a single day on a high-end PC with 192 threads.

## Acknowledgments

This work was supported by NIH HL163585 to CMY, HL163966 to TK, KSM, and KSC, HL136590 to SGC, HL146676 and HL173989 to JES and KSC, HL163977 and NSF 2406028 to JFW, LCL, and KSC, AHA 23TPA1074093 to BCWT, and a Leducq Foundation Network of Excellence to KSC.

## Contributions

John R. Kotter: Conceptualization, data curation, review and editing, validation

Steve Leung: Data curation, review and editing

Thomas Kampourakis: Data curation, review and editing, funding acquisition

Lik-Chuan Lee: Conceptualization, review and editing, funding acquisition

Jonathan Wenk: Conceptualization, review and editing, funding acquisition

Michael Moulton: Conceptualization, data curation, review and editing

Bertrand Tanner: Review and editing, funding acquisition

Stuart Campbell: Conceptualization, review and editing, funding acquisition

Christopher Yengo: Review and editing, funding acquisition

Kerry McDonald: Conceptualization, review and editing, funding acquisition

Julian Stelzer: Data curation, conceptualization, review and editing, funding acquisition

Kenneth S. Campbell: Conceptualization, software development, visualization, original draft, review and editing, funding acquisition

