## Supplementary information for "Hearts may grow concentrically to balance ATP supply and demand and eccentrically to stabilize titin-based stress"

Corresponding author:

Kenneth S. Campbell

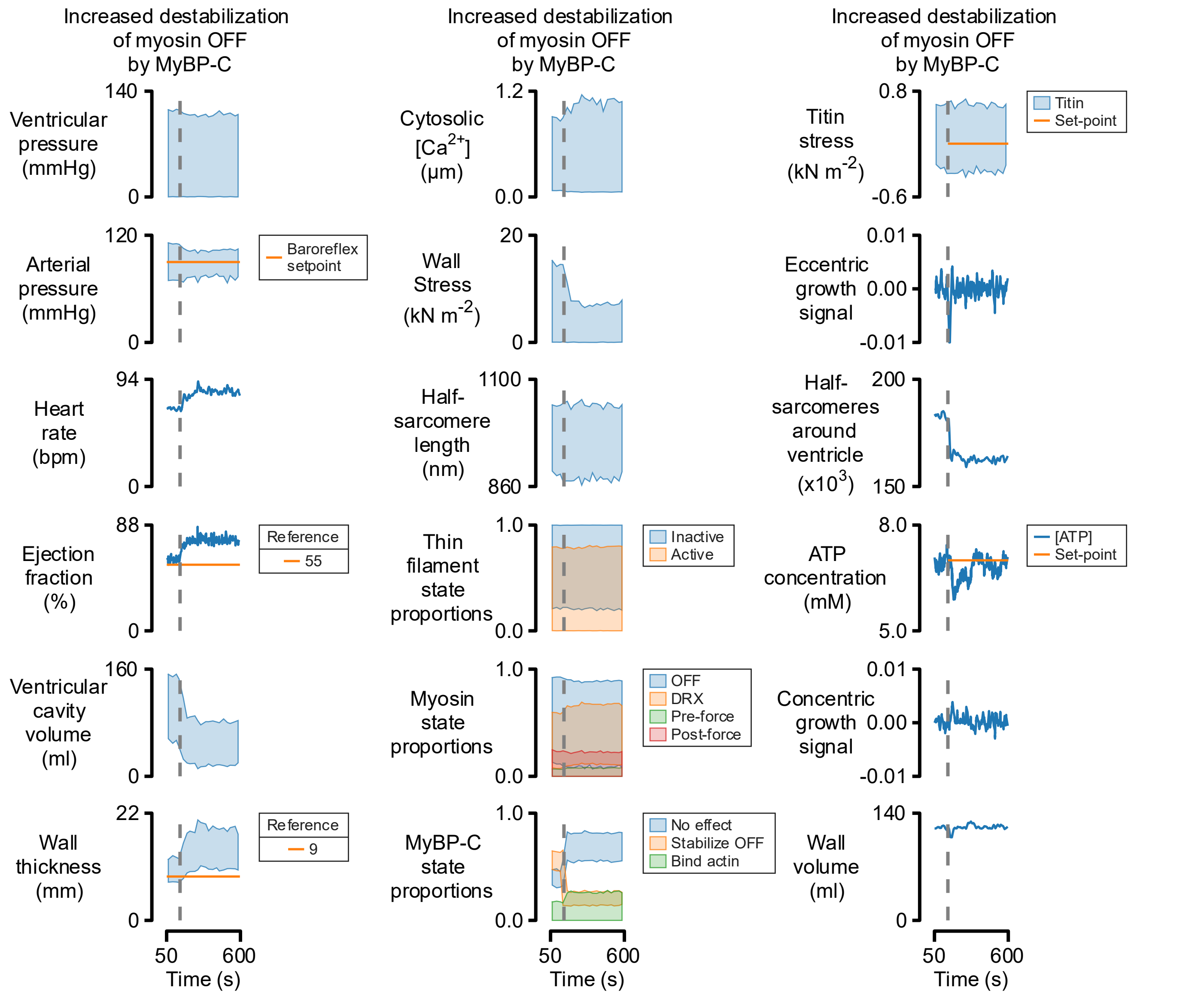


**Figure S1: Destabilization of OFF myosin by a change in the function of myosin binding protein-C (MyBP-C) molecules induces wall thickening and chamber constriction.** Like Fig 2 in the main text, the growth algorithms were activated at 50 s. At 150 s, the function of MyBP-C molecules was adjusted so that they became less effective at stabilizing neighboring myosin molecules in their OFF state.

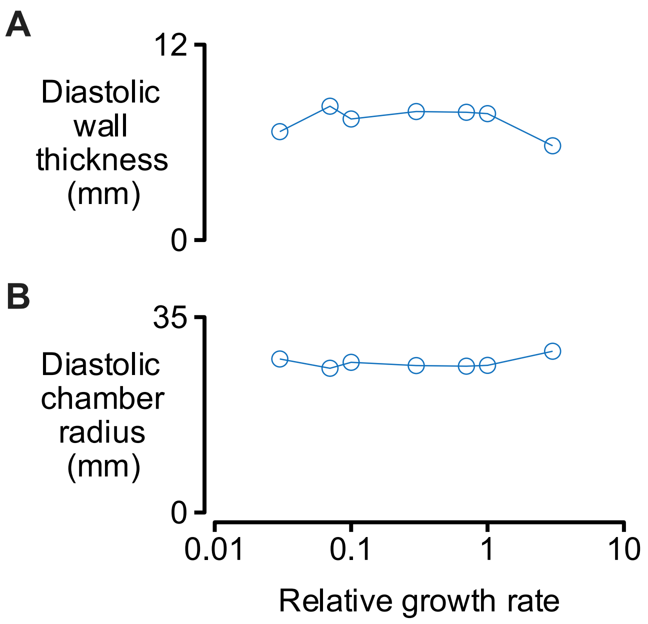


**Figure S2: Slowing growth does not change the predicted dimensions of the ventricle at steady-state.** Data show wall thickness and chamber radius for simulations like those in Fig 2 in the main text implemented with different values of the master growth rate (equations 3 and 6 in main text). If the growth rate was accelerated more than 10-fold, ventricular growth interfered with baroreflex control and the simulations became unstable.

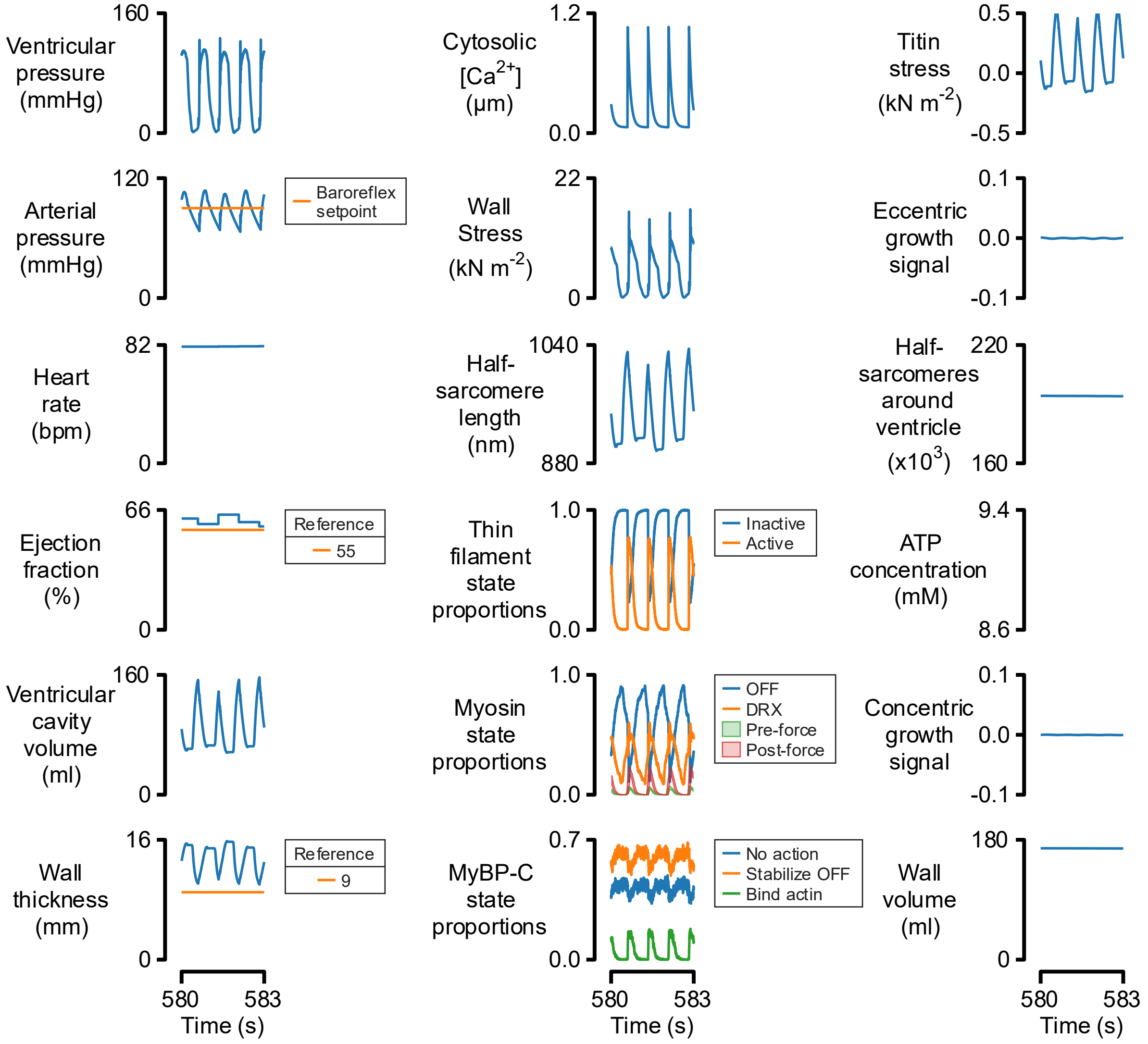


**Figure S3: Predictions for cardiac function during arteriolar dilation.** Heart rate and contractility had increased above basal values to maintain arterial pressure despite a large reduction in arteriolar resistance.

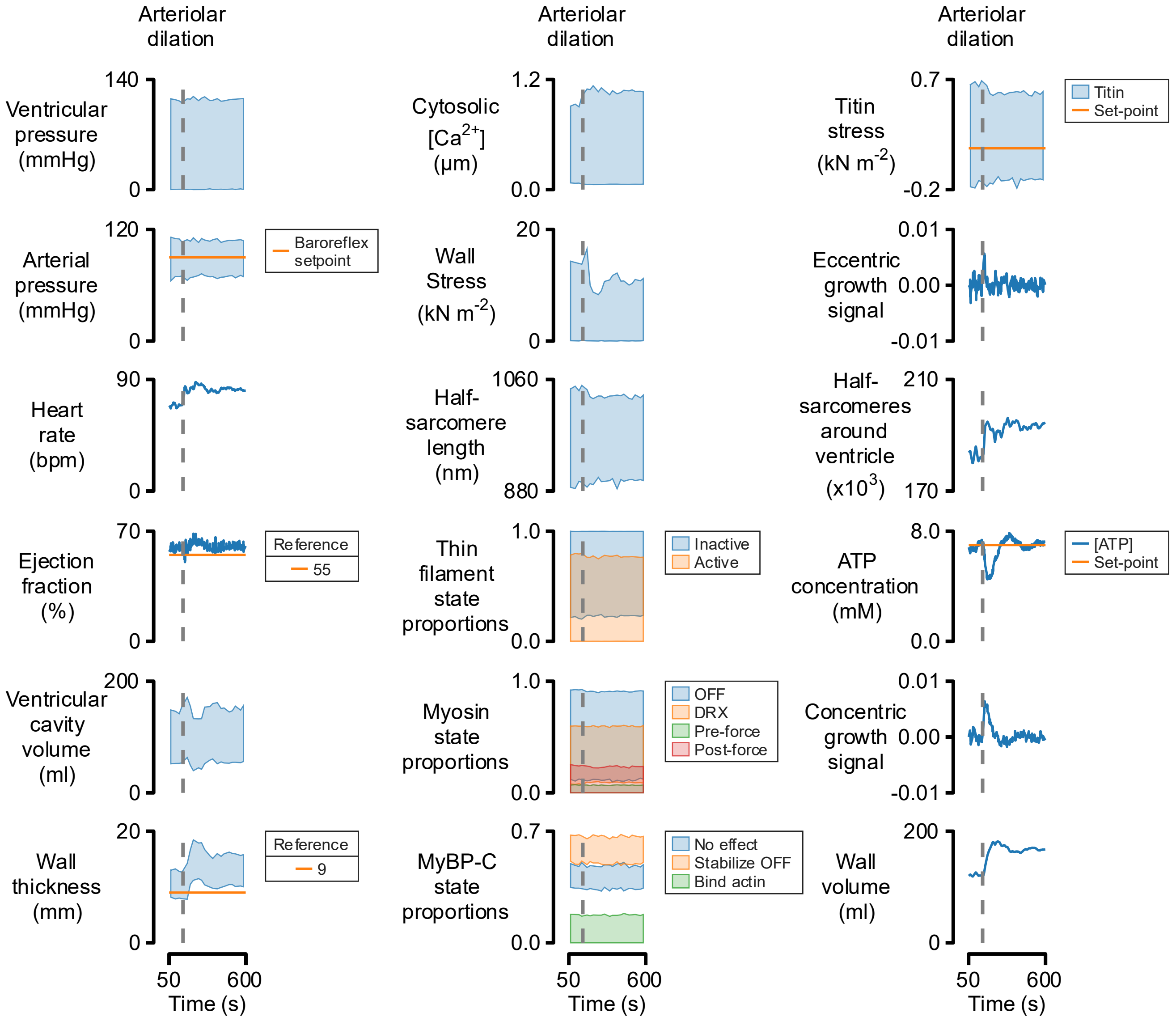


**Figure S4: Arteriolar dilation induces wall thickening.** Similar to Fig 2 in the main text, the growth algorithms were activated at 50 s. At 150 s, the arteriolar resistance was decreased. Autonomic control via the baroreflex increased contractility, heart-rate and vascular tone to maintain arterial pressure. The ventricle thickened to sustain the increased cardiac power.

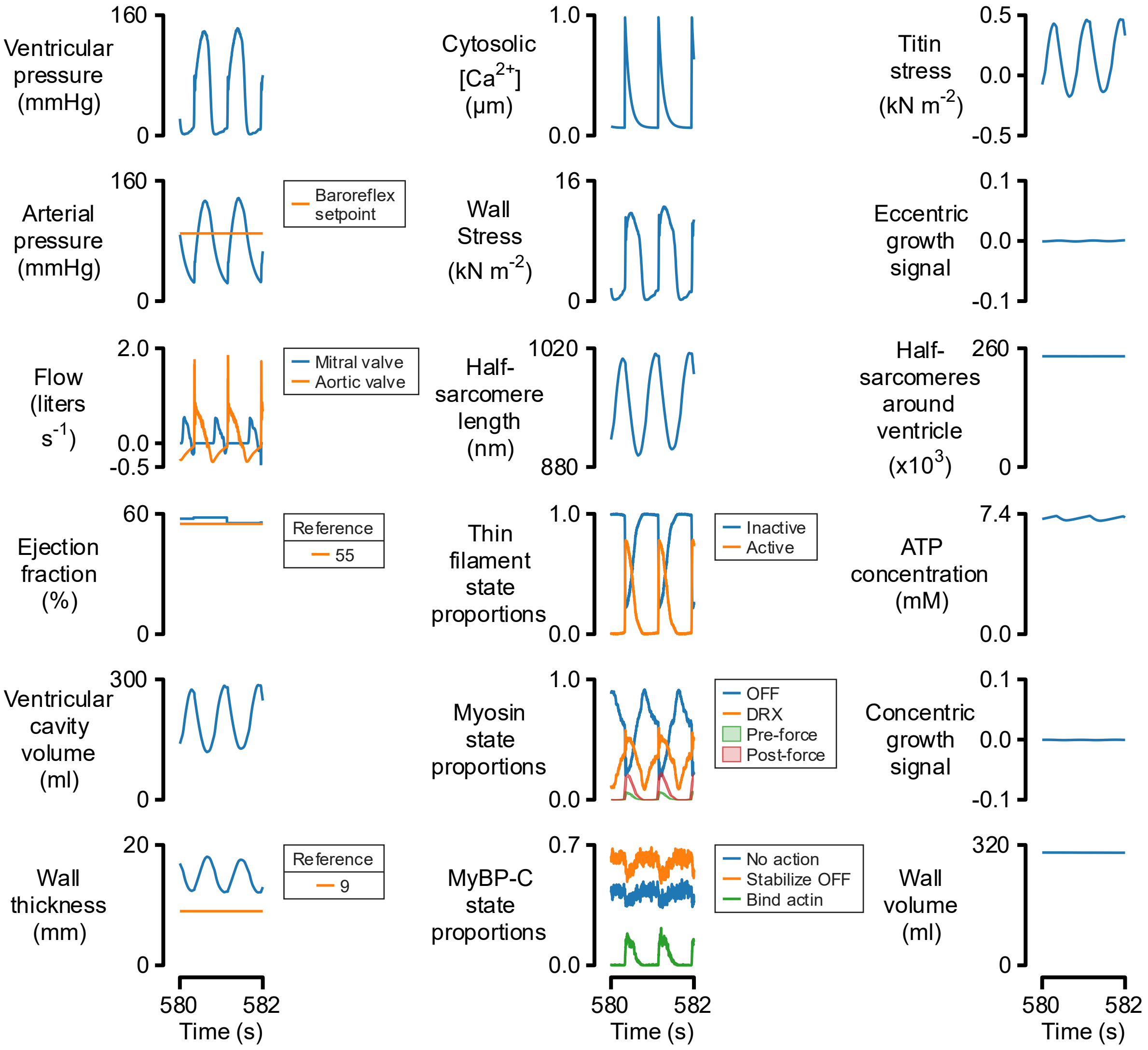


**Figure S5: Predictions for cardiac function in the presence of aortic valve insufficiency.** Note the flows through the aortic and mitral valves in the first column, third row. The sharp transients are associated with the finite time for the valves to open and close in the simulations.

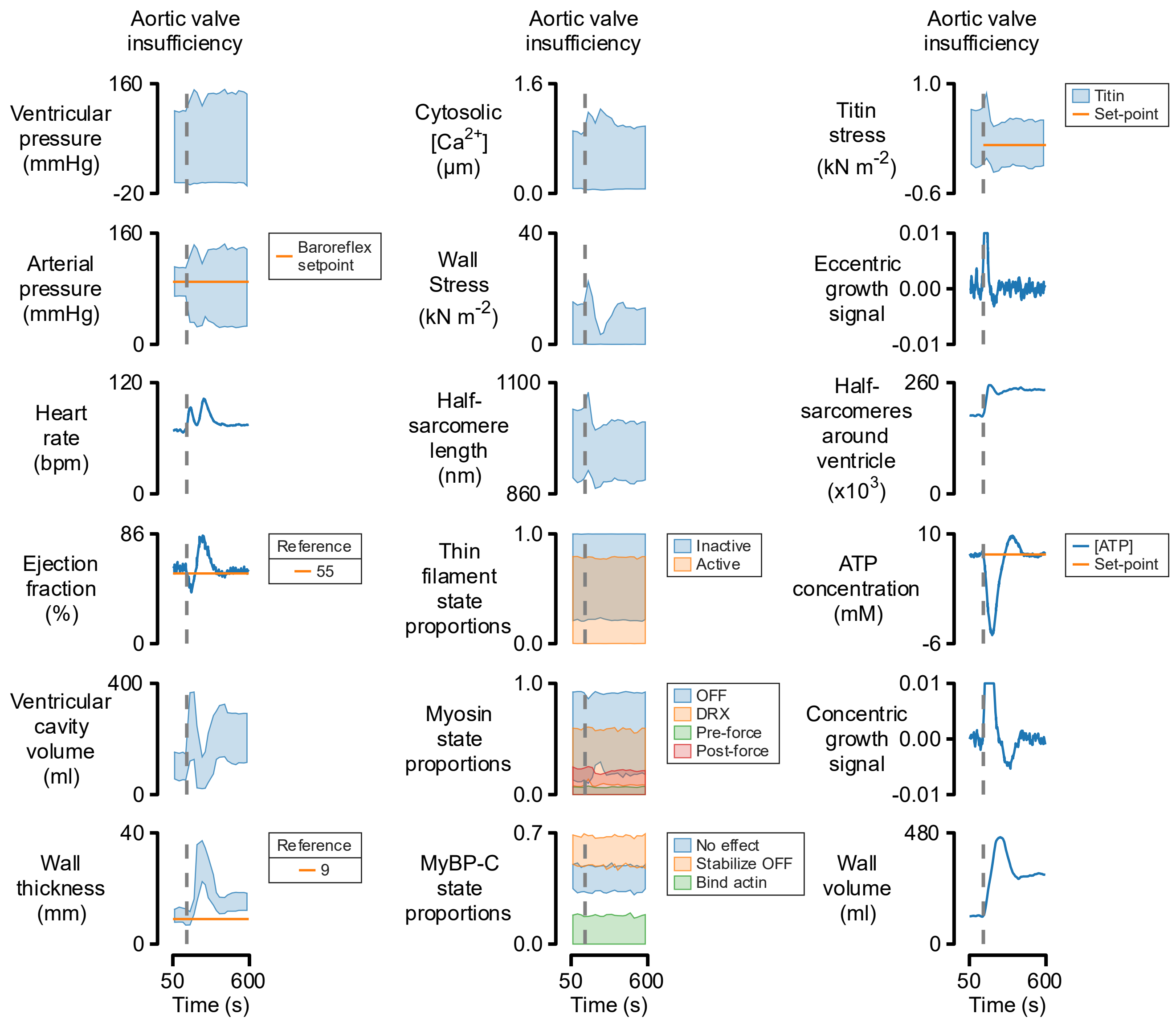


**Figure S6: Aortic valve insufficiency induces wall thickening and chamber dilation.** Similar to Fig 2 in the main text, the growth algorithms were activated at 50 s. At 150 s, the aortic valve was made insufficient so that blood leaked backwards into the ventricle during diastole.

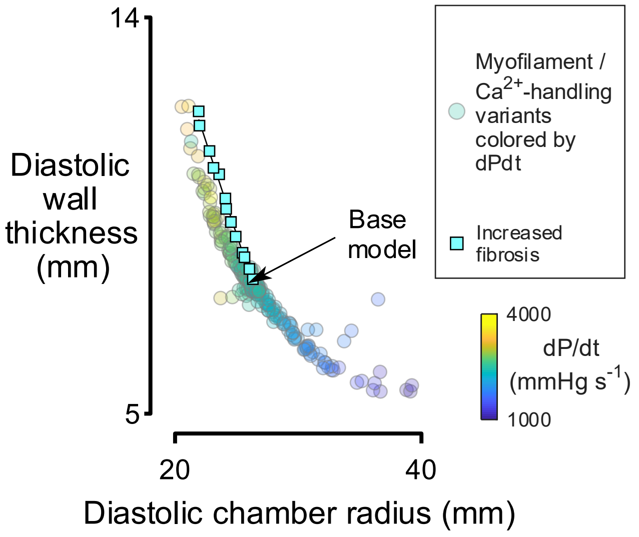


**Figure S7: Increased fibrosis thickens ventricular walls and constricts chambers.** Circles are replotted from Fig 3 in the main text and show the predicted response when myofilament or calcium-handling related properties were adjusted to change contractility. The squares show the responses to progressive increases in the proportion of myocardium occupied by fibrosis.
